# Comparative Phylogenetic Analysis and Transcriptomic Profiling of Dengue (DENV-3 genotype I) Outbreak in 2021 in Bangladesh

**DOI:** 10.1101/2022.10.29.514352

**Authors:** Md. Murshed Hasan Sarkar, M. Shaminur Rahman, M. Rafiul Islam, Arafat Rahman, Md. Shariful Islam, Tanjina Akhtar Banu, Shahina Akter, Barna Goswami, Iffat Jahan, Md Ahashan Habib, Mohammad Mohi Uddin, Md. Ibrahim Miah, Aftab Ali Shaikh, Md. Salim Khan

## Abstract

**Objectives:** The lack of a dengue disease animal model and the complex immune interaction in dengue infection hinders the study of host response and immunopathogenesis. The development of next-generation sequencing (NGS) technology allowed the researchers to study the transcriptomic profiles of human in-depth. We, therefore, implicated phylodynamic and transcriptomic approaches through NGS technology to know the origin of dengue virus (DENV) and their host response in infected patients with dengue fever.

**Methods:** To determine the whole genome sequences of the dengue virus and their transcriptomic profiles, RNA was extracted from the serum samples of 3 healthy, and 21 dengue patients. These samples were custom performed at phylogenetic, phylodynamic, differential express gene (DEG), and gene ontology (GO) using respective bioinformatics tools.

**Results:** The whole genome sequence analysis revealed that the total number of nucleotide ranges on these serum RNA samples were in between 10647 and 10707. Phylogenetic tree analysis showed that these strains were belonged to DENV-3 genotype I. Phylodynamic analysis showed that the 2021 epidemic isolates were clade shifted and maintained as a new clade in compared to 2019 epidemic. Transcriptome analysis mapped a total of 19267 expressed genes. Of them, there were higher expression of genes in dengue-positive samples (n = 17375) with a count of 6953 unique genes in comparison to healthy controls (n = 12314) with a count of 1892 unique genes. A total of 2686 DEGs were identified in a host factor-independent manner in dengue patients with a q-value < 0.05. DESeq2 plot counts function of the top 24 genes with the smallest q-values of differential gene expression of RNA-seq data showed that 11 genes were upregulated, whereas 13 genes were downregulated. GO analysis showed a significant upregulation (p = <0.001) in a process of multicellular organismal, nervous system, sensory perception of chemical stimulus, and G protein-coupled receptor signalling pathways in the dengue patients. However, there were a significant downregulation (p = < 0.001) of intracellular component, cellular anatomical entity, and protein-containing complex in dengue patients. Most importantly, there was significant increase of a classes of immunity protein (Cytokines especially TGF-β1, chemokines, inflammasome, and factors for blood coagulations) in dengue patients in compared to the healthy controls, with increased GO of immune system process. In addition, upregulation of toll receptor (TLR) signalling pathways were also initiated in the patients infected with dengue virus. These TLR pathways were particularly involved for the activation of innate in couple with adaptive immune system that probably involved the rapid elimination of dengue virus infected cells. These differentially expressed genes could be further investigated for target based prophylactic interventions for dengue.

**Conclusion:** This is a first report to document the complete genomic features of dengue, and differentially expressed genes in patients with dengue virus in Bangladesh. These genes may have diagnostic and therapeutic values for dengue infection. Continual genomic surveillance is required to further investigate the shift in dominant genotypes in relation to viral pathogenesis.

## 1. Introduction

The dengue virus is transmitted by mosquitoes and causes a global outbreaks and epidemics in tropical and subtropical areas, with nearly 400 million cases annually, posing an immediate threat to human health in developing countries like Bangladesh (Bhatt et al., 2013). Though first dengue report in Bangladesh was at 1964 but the outbreaks began in 2000 (Mahbubur Rahman et al., 2002) caused by DENV-3 (Aziz et al., 2002) and continuing with circulating serotype/genotype. In 2000-2009 DENV-3 genotype II were prevalent but 2013-2016 DENV-1 and DENV-2 were the most prevalent serotypes in Bangladesh (Muraduzzaman et al., 2018). The serotype DENV-3 was predominant in outbreaks in 2017-2019 and reported to have clad shift to genotype I (Suzuki et al., 2019). The World Health Organization (WHO) reports that 500,000 cases of severe dengue each year require hospitalization, mainly among children (Nikolayeva et al., 2018). Clinically, dengue virus (DENV) diseases can be classified in to two categories, uncomplicated dengue or dengue fever (DF), dengue hemorrhagic fever (DHF), or dengue with warning signs (Bhatt et al., 2013). A hallmark of DHF is increased vascular permeability, resulting in plasma leakage, rash, bleeding, and cardiovascular collapse. The leading cause of death and morbidity in DHF is vascular leakage and its secondary complications (Saini et al., 2020). DENV consists of four genetically associated serotypes (DENV-1 to DENV-4), which can be further subdivided into genotypes (Holmes & Twiddy, 2003). Infection with one type of dengue virus confers life-long immunity; subsequent infections with other types of the virus do not confer immunity (Mukhopadhyay et al., 2005). The shift of a predominantly circulating dengue serotype and/or genotype is responsible for increasing the incidence and severity of dengue outbreaks worldwide (Nunes et al., 2016). Dengue serotyping/genotyping shifting facilitate infectivity via a non-neutralizing cross-reactive antibody that enhance disease by binds to the second infecting DENV virus serotype result of accelerate virus entry into cells through an Fc receptor-mediated endocytosis and by suppressing intracellular innate responses against the virus (Bournazos et al., 2020; Chareonsirisuthigul et al., 2007). Similarly, DENV vaccine recipients who have previously been exposed to DENV may be at risk for serious illness because of the increased virus production and suppressed antiviral defenses (Deng et al., 2020). Live attenuated vaccines elicit both the innate immune system and provide antigens to stimulate adaptive immunity (Iwasaki & Medzhitov, 2015). DENV infection triggers an innate immune response, which alters gene expression profiles and leads to host-pathogen interactions (M. J. Li et al., 2020). This interaction is mediated by several pattern-recognition receptors (PRRs), including the endosome-associated Toll-like receptor-3 (TLR-3) and TLR-7 and the cytoplasmic retinoic acid-inducible gene I (RIG-I), NOD-like receptor protein 3 (NLRP-3)-specific inflammasome and melanoma differentiation-associated protein 5 (MDA-5), are responsible for activating downstream signaling pathways via recognition of their ligands (Medzhitov, 2007). Activating downstream signaling pathways can result in the production of type I and type III interferons (IFNs) and proinflammatory cytokines, and hundreds of interferon-stimulated genes (ISGs) that help prevent DENV infection (Hur, 2019; Lazear et al., 2019). IFN production is intrinsically linked to the transcriptional and posttranslational modification and regulate gene expression in the signaling pathways that stimulate DENV-specific adaptive immunity (Carpenter et al., 2013). A number of classical innate immunity pathways, such as RNAi, TLR, and JAK/STAT pathways, can inhibit viral infection and immunity in mosquitoes (Souza-Neto et al., 2009). Dengue virus inhibits IFN-α and IFN-β signaling by suppressing the JAK-STAT pathway, resulting in reduced host immunity against the virus (Muñoz-Jordán et al., 2003). However, DENV infections alter gene expression profiles in dengue infected patients.

The combination of several factors such as reduced host defense, increased virus uptake and delayed viral clearance show synergism to produce higher virus titer that causes severe outcome. Early identification is more challenging due to the diverse clinical manifestation of dengue infection, as well as its substantial similarities to other febrile viral infections (Saini et al., 2020). To control the dengue diseases, it is necessary to identify the dengue serotype, genotype, diseases process, pathogenesis, and host response to dengue infection. The genomic characteristic of DENV outbreak and host response to dengue infection in Bangladesh is largely unknown. So we have therefore chosen to study the genomic characteristic and host response of DENV infection.

The study of host response and immunopathogenesis in dengue infection is challenging due to the lack of disease model in animals and the complex immune interaction with the DENV infection. The development of high-throughput sequencing technology radically changes the approaches for understanding and analyzing the transcription level information between host virus interactions. This advance technology helps to design novel strategies for blocking virus transmission and detection marker for diagnosis. A number of research areas have used this technology as a tool to generate hypothesis including the identification of drug action mechanisms, cellular responses, and drug target discovery (Kobasa et al., 2007).

In this study we extensively analyze the transcriptome as well as alternatively spliced transcriptome of blood samples collected from participants with and without dengue infection. We also reveal the whole genome sequence (WGS) of dengue viruses from our samples to determine serotype, genotype, phylogenetic relation and phylodinamics. Based on the transcriptomic signatures elicited by dengue patients, we demonstrate that differential expression patterns of gene expression and transcript isoform expressions, as well as differential splicing patterns, contribute to defining early transcriptomic signatures of diagnostic values. Gene ontology (GO) analysis depicts the function of the differentially expressed gene.

Our data indicate that signatures of innate immunity correlate with the dengue infection. It’s also shown that dengue infection up-regulate the biological adhesion, blood coagulation process though these are the characteristic features of DHF. TGF-1 was a highly activated upstream regulator in the positive sample. Collectively, these studies demonstrate specific transcriptional pathways and immune cell types implicated in inducing IFNs and ISGs, emphasizing the importance of gene regulation mechanisms at the splicing and isoform levels for eliciting an immunological response. Our study shows for the first time that a comprehensive view of transcriptome expression in plasma of dengue patients in Bangladesh. It offers a valuable resource for understanding transcriptome that might be involved in the manifestation and progression of dengue. It might help to discover valuable gene for diagnosis, therapeutic and key transcriptional regulatory factors that will block viral replication and transmission in mosquitoes. We also clearly showed that clade shifting which might be a predictor of dengue severity.

## 2. Methodology

### a. Sample Collection

Blood sample were collected in October 2021 from febrile patients who were clinically suspected to have dengue by the Public Health Authority for diagnosis of dengue. To detect dengue the NS1 antigen (NS1 present from the first day of the disease) was first detected using the NS1 Ag rapid assay kit following the manufacturer’s instruction. Serum was separated from the blood and RNA was extracted from 200 μL serum sample of each aliquot with the ReliaPrep™ Viral TNA Miniprep System (Promega), according to the manufacturer’s protocol. The extracted RNAs were freshly used for laboratory tests or stored at −80°C. A total of 24 samples were obtained, of which 21 were positive for dengue and 3 were from healthy persons whose NS1 serum tests were negative.

### b. Library Preparation and Sequencing

Viral cDNA and DNA libraries were construct using Illumina TruSeq RNA Library Prep Kit (Illumina) following the manufacturer’s instruction. After libraries preparation, the samples were sequenced using the NextSeq 550 sequencing System with an output of paired◻end (2 × 74 bp) reads.

### c. Whole genome assembly and genotyping

The Illumina sequencer creates raw pictures by employing sequencing control software for system control and base calling via RTA, which is an integrated primary analysis program (Real Time Analysis). The binary BCL (base calls) is translated into FASTQ using the Illumina software bcl2fastq (https://support.illumina.com/sequencing/sequencing_software/bcl2fastq-conversion-software.html), resulting in an average of ~2.61 million reads per sample. FastQC v0.11 was used to assess the quality of the produced FASTQ files (Andrews & others, 2010). Trimmomatic v0.39 (Bolger et al., 2014) was used to clip adapter sequences and low-quality ends per read with predefined parameters of sliding window size 4; a minimum average quality score of 20; and a minimum read length of 36 bp. After trimming, an average ~2.53 million reads per sample passed the quality testing procedures (Table 1). By aligning a reference genome (GenBank accession no. NC_001475.2), the consensus was obtained using the Burrows-Wheeler Aligner (BWA v0.7.17) (H. Li & Durbin, 2009), SAMtools v1.12 (H. Li et al., 2009), and BEDTools v2.30.0 (Quinlan & Hall, 2010). Snippy was used to assess indel and area-wise mutation coverage, and the genome was repaired accordingly. Freebays (https://github.com/freebayes/freebayes) is the variation caller of snippy, with the minimum number of reads covering a site to be considered (default=10) and the minimum VCF variant call “quality” (default=100) (Seemann, 2015). GENOME DETECTIVE VIRUS TOOL (https://www.genomedetective.com/app/typingtool/virus/) was also implemented to virus detection and *de-novo* assembly of the virus (Vilsker et al., 2019). Genome Detective begins by categorizing short reads into groupings, or buckets. All readings from a single viral species are placed in the same bucket. DIAMOND (Buchfink et al., 2014) is then allocated to each bucket for taxonomic identification. Once all of the readings have been sorted into buckets, each bucket is constructed from scratch using metaSPAdes (Nurk et al., 2017). BLASTx and BLASTn employ contigs to search for possible reference sequences against the National Center for Biotechnology Information (NCBI) RefSeq viral database. Advanced Genome Aligner (AGA) (Deforche, 2017) is used to link the contigs for each particular species (Deforche, 2017). A report is created that refers to the final contigs and consensus sequences, both of which are accessible as FASTA files (Vilsker et al., 2019). DENGUE VIRUS TYPING TOOL (https://www.genomedetective.com/app/typingtool/dengue/) implemented in Genome detective (Vilsker et al., 2019) were used to identify virus genotype.

**Table 1:** Sample and sequence information of the collected samples in this study.

| Sample Name | Sex | Age | Total read | Read after QC | Length | Percent covered (%) | Mean coverage (×) | Genotype | GenBank Accession No. |
| --- | --- | --- | --- | --- | --- | --- | --- | --- | --- |
| BCSIR-05 | Female | 26 | 4311927 | 4227543 (98.04%) | 10707 | 98.89 | 28.15 | DENV-3 Genotype I | SAMN21369031 |
| BCSIR-IBS-D1 | Male | 12 | 3507870 | 3452844 (98.43%) | 10686 | 99.87 | 48375.9 | DENV-3 Genotype I | ON103303 |
| BCSIR-IBS-D2 | Female | 30 | 3673245 | 3618733 (99.00%) | 10686 | 99.73 | 16465.9 | DENV-3 Genotype I | ON103391 |
| BCSIR-IBS-D3 | Female | 8 | 3249713 | 3143738 (96.74%) | 10672 | 99.57 | 3415.74 | DENV-3 Genotype I | ON111446 |
| BCSIR-IBS-D4 | Male | 61 | 4200439 | 4120093 (98.09%) | 10703 | 99.88 | 82474.8 | DENV-3 Genotype I | ON115022 |
| BCSIR-IBS-D5 | Male | 21 | 2428255 | 2150805 (88.57%) | 10669 | 99.64 | 11612.4 | DENV-3 Genotype I | ON115024 |
| BCSIR-IBS-D6 | Male | 4 | 2476067 | 2454059 (99.11%) | 10667 | 99.55 | 2491.9 | DENV-3 Genotype I | ON115026 |
| BCSIR-IBS-D7 | Male | 31 | 3347486 | 3291974 (98.34%) | 10683 | 99.71 | 3882.28 | DENV-3 Genotype I | ON115030 |
| BCSIR-IBS-D8 | Male | 10 | 3585995 | 3564733 (99.41%) | 10688 | 99.89 | 72808.2 | DENV-3 Genotype I | ON115183 |
| BCSIR-IBS-D9 | Female | 3 | 1372523 | 1196648 (87.19%) | 10669 | 99.51 | 708.61 | DENV-3 Genotype I | ON115812 |
| BCSIR-IBS-D10 | Female | 41 | 3921170 | 3885916 (99.10%) | 10686 | 99.93 | 68242.7 | DENV-3 Genotype I | ON115815 |
| BCSIR-IBS-D11 | Male | 24 | 2418634 | 2372894 (98.11%) | 10660 | 97.78 | 140.457 | DENV-3 Genotype I | ON115817 |
| BCSIR-IBS-D12 | Male | 26 | 2329130 | 2288986 (98.28%) | 10667 | 99.51 | 4856.02 | DENV-3 Genotype I | ON115819 |
| BCSIR-IBS-D13 | Female | 60 | 2782261 | 2631028 (94.56%) | 10668 | 99.20 | 401.65 | DENV-3 Genotype I | ON116130 |
| BCSIR-IBS-D14 | Female | 22 | 2758002 | 2688567 (97.48%) | 10707 | 99.74 | 26583.9 | DENV-3 Genotype I | ON127419 |
| BCSIR-IBS-D15 | Male | 37 | 1743673 | 1712426 (98.21%) | 10707 | 99.56 | 5517.04 | DENV-3 Genotype I | ON127534 |
| BCSIR-IBS-D16 | Male | 50 | 2964830 | 2913773 (98.28%) | 10660 | 98.26 | 132.21 | DENV-3 Genotype I | SAMN31519327 |
| BCSIR-IBS-D17 | Male | 12 | 2008835 | 1990945 (99.11%) | 10669 | 99.56 | 10660.1 | DENV-3 Genotype I | ON127538 |
| BCSIR-IBS-D18 | Male | 24 | 2370633 | 2314886 (97.65%) | 10707 | 99.73 | 14478.9 | DENV-3 Genotype I | ON127551 |
| BCSIR-IBS-D19 | Female | 5 | 310019 | 300836 (97.04%) | 10647 | 99.44 | 87.73 | DENV-3 Genotype I | ON127553 |
| BCSIR-IBS-D20 | Male | 50 | 2452702 | 2403740 (98.00%) | 10686 | 99.05 | 129.598 | DENV-3 Genotype I | SAMN31519328 |
| BCSIR-IBS-D21 | Male | 35 | 1543653 | 1445717 (93.66%) | - | - | - | - | - |
| BCSIR-IBS-D22 | Male | 52 | 1369461 | 1172879 (85.65%) | - | - | - | - | - |
| BCSIR-IBS-D23 | Male | 55 | 1419348 | 1348541 (95.01%) | - | - | - | - | - |

### d. Phylogenetic analysis

NCBI BLASTn was used to find the most closely related strain, and representative sequences were retrieved for phylogeny. Outgroup DENV-3 genotypes II, III, and V were also collected, and all sequences were genotyped using the DENGUE VIRUS TYPING TOOL (https://www.genomedetective.com/app/typingtool/dengue/). Multiple alignment using fast Fourier transform (MAFFT) online version (v7.0) (https://mafft.cbrc.jp/alignment/server/) were used for multiple sequence alignment with their default parameters (Katoh & Standley, 2013). The complete viral genome was built using a neighbor joining (NJ) phylogenetic tree (Saitou & Nei, 1987) with a bootstrap value of 1000 and Jukes-Cantor substitution model (Erickson, 2010). Phylo.io (http://phylo.io/) was used to visualized the phylogenetic tree.

Another *env* gene sequence based phylogenetic tree was reconstructed that has more DENV3 isolates reported from Bangladesh (Suzuki et al., 2019) which contains 135 isolates. After making ClustalW alignment, a model selection was carried out based on the Akaike information criterion in MEGA11 (Kumar et al., 2012). A neighbor-joining tree was reconstructed using the Tamura-Nei model with Gamma distributed rates among sites. 1000 bootstrap replications were used to test the phylogeny.

### e. Phylodynamic analysis

Bayesian inference (BI) analyses were conducted by using Timra-Nei with a gamma-distributed rate variation substitution model (TN93 + G). The Markov Chain Monte Carlo (MCMC) algorithm in the BEAST package v.1.10.4 was used (Drummond & Rambaut, 2007). The calibration point was the date of isolation of each sample. Runs were performed using the Bayesian skyline as the tree prior under the uncorrelated relaxed molecular clock (Ho & Shapiro, 2011). The evolutionary analysis was run for 100 million steps and the trees were sampled every 1,000 states. Convergence of the MCMC chains were inspected using TRACER v.1.7.2 (http://tree.bio.ed.ac.uk) (Rambaut et al., 2018). Posterior trees were summarized by discarding the first 10% of the sampled trees and choosing the Maximum Clade Credibility (MCC) were summarized using TreeAnnotator v.1.10.4. The final tree was then visualized and plotted using FigTree v.1.4.4 (http://tree.bio.ed.ac.uk). A posterior mean for each evolutionary parameter was presented along with a 95% Bayesian credibility interval.

### f. Recombination analysis

The DENV-3 recombination was detected using seven approaches, including RDP (Darren P. Martin et al., 2015), Chimaera (Posada & Crandall, 2001), BootScan (D. P. Martin et al., 2005), 3Seq (Boni et al., 2007), NCONV (Padidam et al., 1999), MaxChi (Smith, 1992), and SiScan (Gibbs et al., 2000), all of which are accessible in the Recombination Detection Program (RDP version 5) (http://web.cbio.uct.ac.za/~darren/rdp.html) (Darren P. Martin et al., 2021), with p < 0.01. If a sequence was discovered by at least three ways using the multiple comparison correction setting option, the recombinant event was identified (Zeng et al., 2018).

### g. Human gene expression analysis

RNA-Seq Alignment tools from illumine basespase (https://www.illumina.com/products/by-type/informatics-products/basespace-sequence-hub/apps/rna-seq-alignment.html) were applied for human gene expression. Here, sample reads were mapped by using the quick universal RNA-seq aligner, STAR (Dobin et al., 2013). The aligner utilized the NCBI GRCh38 human genome as a reference genome. Salmon, a rapid and bias-aware transcript expression quantification tool (Patro et al., 2017) was used to quantification of expressed genes from transcriptome samples.

### h. Statistical analysis

DESeq2 (release 3.13) in R (version 4.0) was used for differential gene expression and statistical analysis. DESeq2 (Love et al., 2014) was chosen since it is a popular parametric tool with a detailed and constantly updated user guideline. (http://bioconductor.org/packages/release/bioc/vignettes/DESeq2/inst/doc/DESeq2.html). Because DESeq2 inherently corrects for library size, it is critical to supply un-normalized raw read counts as input. To adjust for discrepancies in sequencing depth, we utilized a variance stabilizing transformation (Chu et al., 2017). The Benjamini-Hochberg technique (Benjamini & Hochberg, 1995) was used to modify P-values for multiple testing. For the selection of DE genes, a false discovery rate (FDR) adjusted p-value < 0.05 was used. Ggplot2 R package was used to visualize the data. EnhancedVolcano R package was also used for the volcano plot. The data was shown using the R tool ggplot2 (Wickham, 2011). For volcano plotting, the EnhancedVolcano R package was employed https://bioconductor.org/packages/devel/bioc/vignettes/EnhancedVolcano/inst/doc/EnhancedVolcano.html and pheatmap package was employed for heatmap (Kolde, 2015).

### i. Gene ontology (GO) and pathway analysis

PANTHER Functional classification viewed in gene list and Overrepresentation Test (published on December 01, 2020) in PANTHER version 16.0 (http://www.pantherdb.org/) were used to undertake gene ontology and PANTHER pathway studies (Mi et al., 2021). For statistical analysis, PANTHER employed the Fisher Exact test and the False Discovery Rate with the default parameters. Only pathways and GO terms with an FDR p-value < 0.05 were reported in this study.

## 3. Results

Dengue is a one of the most common mosquito-borne viral diseases that affect millions of people each year globally. WHO state that Asia is responsible for 75% of all dengue cases worldwide, especially in countries such as the Philippines, Indonesia, and Thailand. The Dengue seroprevalence in Bangladesh is lower than that of the Southeast Asian countries, but the trend is changing rapidly. Bangladesh experienced its largest dengue outbreak on record during the year 2019 (Lancet). However, there is a lack of genomic surveillance of Dengue in the country. As a result, in this study, we compared RNA-seq data from dengue patients to examine the whole genome sequence, genotype, and genomic characterization of the dengue virus, as well as differential gene expression, overrepresented pathways, and upstream regulators.

This study includes a total of 21 dengue patients and 3 healthy controls. Both male (n = 13) and female (n = 8) patients with ages ranging from 3 to 61 years were enrolled in the study (Table 1). The average days of symptoms were 4.5. At the time of patient recruitment, a blood sample was taken.

### 3.1 Metagenomic sequencing and annotation of the dengue virus genomes

The complete viral genomes of 21 samples positive for Dengue virus were assembled by metagenomic sequencing. The average coverage of the assembled genomes ranged from 28.15 × to 82474.8 × and the genome assembly length varied between 10,647 bp (sample: BCSIR-IBS-D19) and 10,707 bp (sample: BCSIR-05) (Table 1). All the positive samples (n = 21) belonged to DENV-3 Genotype I (Table 1, Figure 1). No reads from the healthy control (n = 3) aligned with the Dengue virus (Table 1).

**Figure 1.**
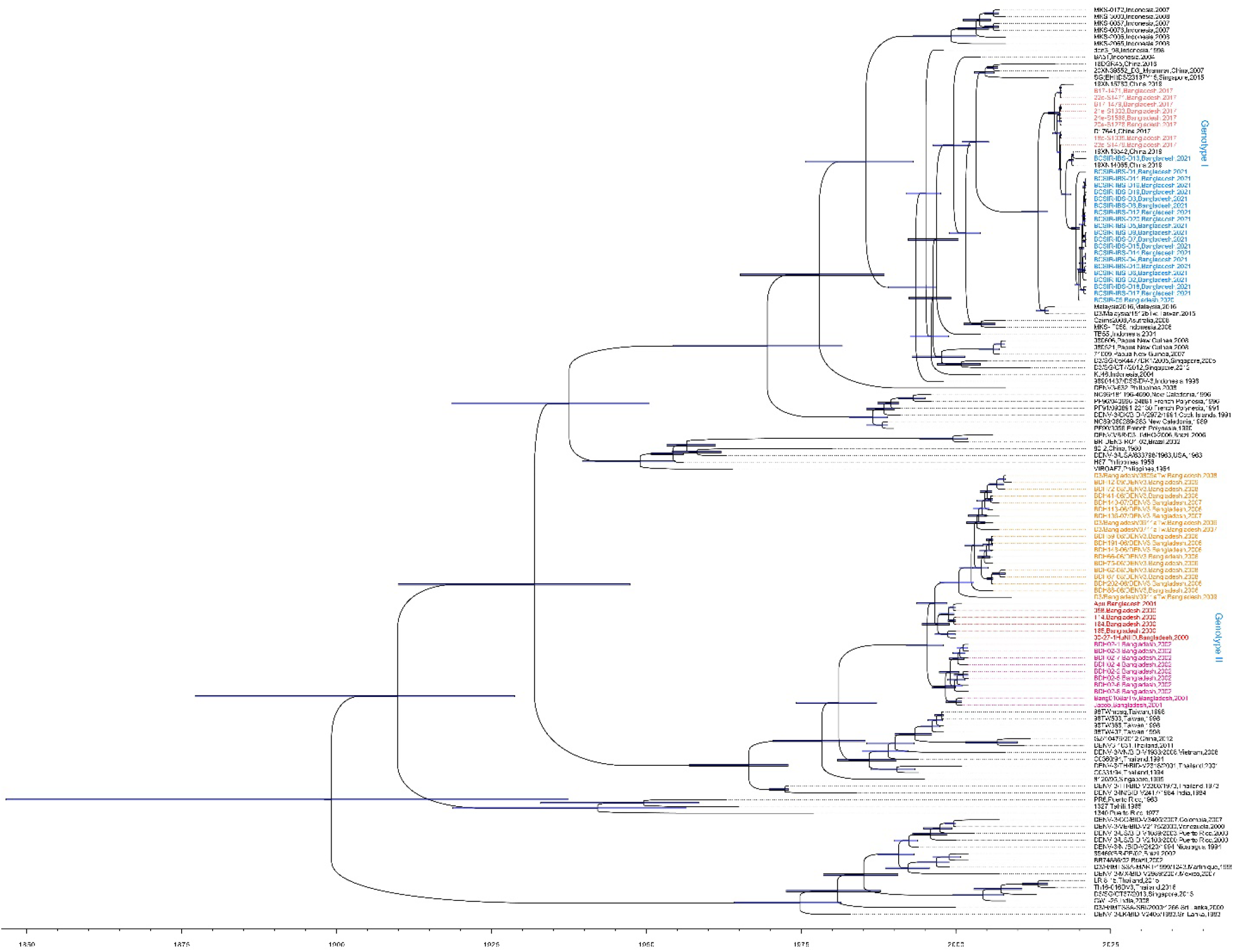
Phylodynamic analysis showed clad shift in DENV3 epidemic in Bangladesh. Bayesian Inference analysis compared all the sequences used to construct a phylogenetic tree. Bars in the internal nodes indicates 95% HPD intervals. Bangladeshi DENV-3 sequences distributed in two clades characterized as genotype I for DENV3 epidemic of 2017 and 2021, and the previous epidemics in genotype II that contains isolates from 2009, 2008, 2006, 2002, and 2000. The neighbor-joining tree was reconstructed using Tamura-Nei model with Gamma distributed rates among sites. 1000 bootstrap replications were used to test the phylogeny.

### 3.2 Phylogenetics and phylodynamics of dengue virus in Bangladesh

After analyzing these sequences, we investigated their phylogenetic relationships to existing dengue virus genomes based on both the complete genomes and the envelope protein E. In both phylogenetic trees, the genomes from this study belonged to DENV-3 Genotype I and the genomes of these samples were highly similar and belonged to a cluster of samples of DENV-3 genotype I originated from China and Thailand that also includes some sequences from Bangladesh collected between 2009 and 2017 (Figure 2.A). In both the trees, isolates from Genotype II, III, and I were clustered separately. The *env* gene tree has more DENV-3 isolates from the previously reported Dengue epidemic in Bangladesh (Supplementary figure 1). It shows that 2021 epidemic samples are closely related to 2017 Dengue epidemic in Bangladesh.

**Figure 2.**
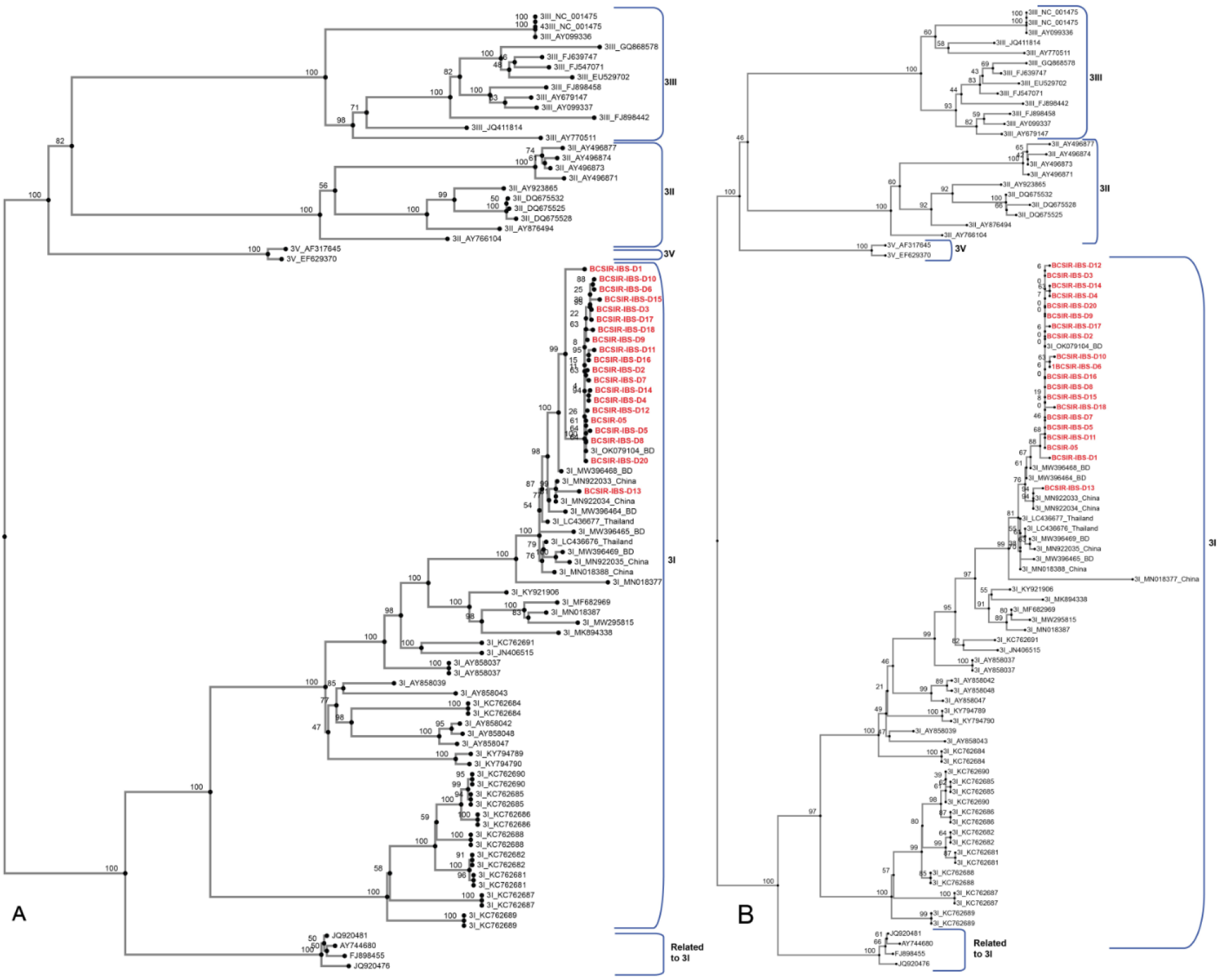
Neighbor joining (NJ) phylogenetic tree of dengue whole genome and envelope protein E. (A) A neighbor joining (NJ) phylogenetic tree with a bootstrap value of 1000 and the Jukes-Cantor substitution model were used to construct the whole viral genome and (B) the envelope protein E phylogenetic tree of 95 DENV-3 strains. The DENV-3 genotype I strain sequenced in this study is shown in red. To root the tree, the full sequences of DENV-3 genotypes II, III, and V were employed.

Bayesian inference analysis comparing all the sequences produced a phylogenetic tree that has similar topography to the previously described NJ tree. The Bangladeshi DENV-3 sequences are distributed in two clades (Figure.1), characterized as genotype I for DENV3 epidemic of 2017 and 2021, and the previous epidemics in genotype II that contains isolates from 2009, 2008, 2006, 2002, and 2000. This confirms a previously reported clad shift in DENV3 epidemic in Bangladesh (Suzuki et al., 2019). However, the tree also indicates that the 2021 epidemic isolates, which have strong similarities among themselves, are well separated from 2017 epidemic isolates in Bangladesh but interleaved by two DENV3 isolates sampled in 2019. These two isolates, 19XN13542 and 19XN14065, were sampled from travelers return from Bangladesh based on the NCBI GenBank description. This gives us an observational time point between 2017 and 2021 and indicates that the DENV3 was evolving in Bangladesh in-between times. The estimated median tMRCA from genotype I DENV3 sequences from Bangladesh (epidemic of 2017 and 2021) was 2013 (95% HPD 2010 – 2014). This was right after the 2006-2009 DENV3 occurrence from genotype II reported in the previous study, where previously reported clad-based genotype shift presumably happened.

### 3.3 Principal component analysis (PCA) and hierarchical clustering analysis

We performed PCA to investigate the clustering of the samples in each group (patients and healthy); whether they clustered within the same group or with other groups. First, we used HTSeq (Anders et al., 2015) to count reads that uniquely aligned to one gene, and PCA plots were then generated by importing the data into DESeq2 (Love et al., 2014). The PCA results demonstrated that most of the positive samples clustered together distinctly separated from the samples of the control group (Figure 3). However, the control group did not cluster so closely, likely due to lower sample number (n = 3) in the control group. Additionally, PCA scree plots showed that principal component 1 (PC1) and 2 (PC2) accounted for 36% and 12%, respectively, of the analyzed data (Figure 3).

**Figure 3.**
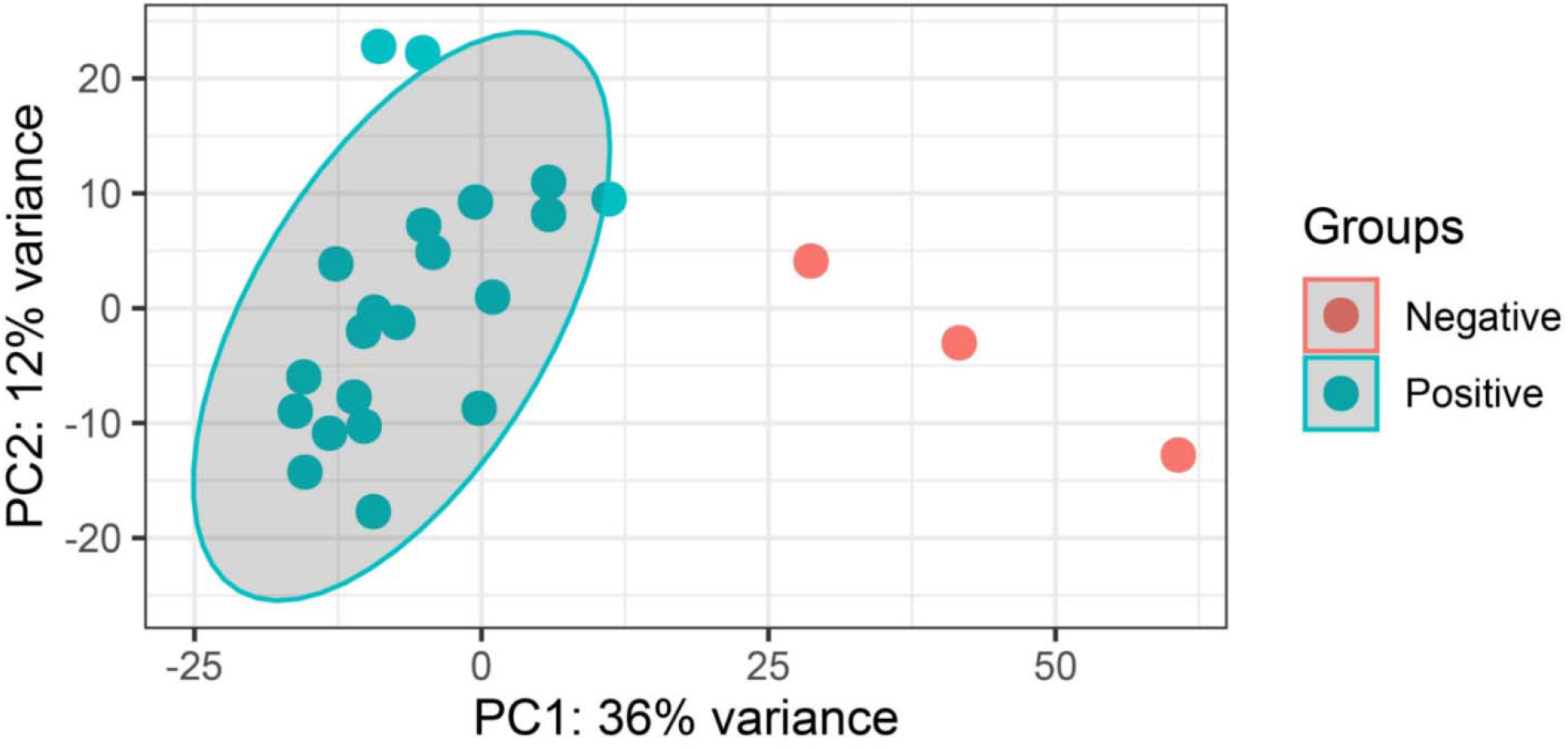
Dengue positive and negative groups were subjected to principal component analysis (PCA). The variance stabilizing transformation in DESeq2 was used to identify principal component 1 (PC1) and principal component 2 (PC2). The percentage of variance represents the amount of variance explained by PC1 and PC2.

### 3.4 Differentially expressed genes (DEGs)

A total of 19,267 genes were identified by at least one read from all the samples including dengue positive and negative (control) samples. Positive samples aligned to a total of 17,375 genes whereas a total of 12,314 genes aligned in the negative control samples. Among them, 6,953 and 1,892 unique genes found in dengue-positive and healthy control samples respectively (Figure 4, Supplementary Data 1). Overall, 2,686 differentially expressed genes (DEGs), with a q-value < 0.05, were detected in the analysis of DESeq2 (Supplementary Data 1).

**Figure 4.**
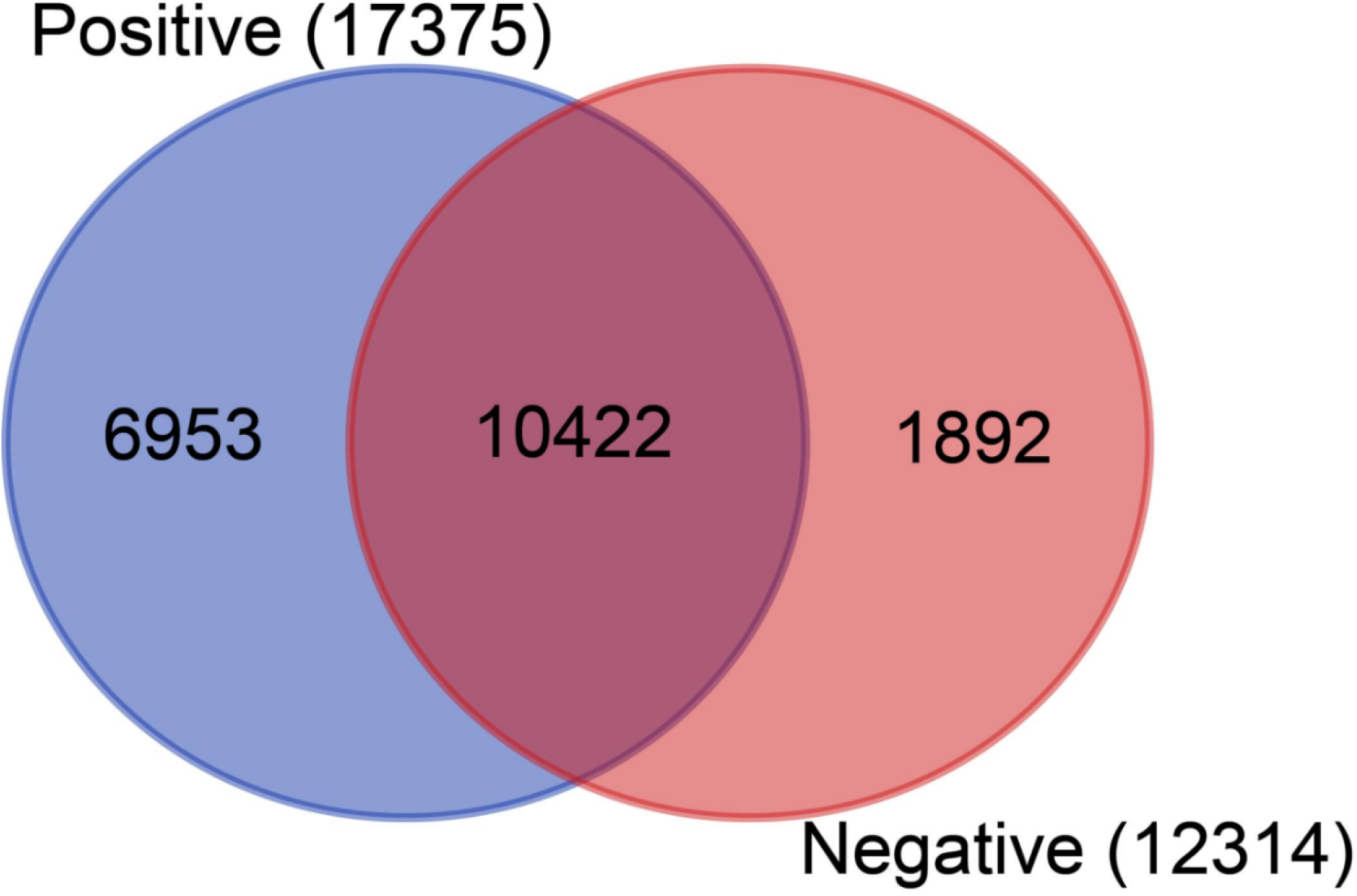
Venn diagram of transcriptome response of dengue positive and negative groups. Venn diagram represents the number of unique and shared genes between dengue positive and negative patients. The total number of genes discovered in each category is shown in parentheses.

The genes were further analyzed using hierarchical clustering and heatmap, with < 25 genes filtered off every row. In the heatmap, positive and control groups created distinct clusters, while 7 positive samples (BCSIR-IBS-D9, D5, D8, D19, D4, D1, D10) sub-clustered from the rest of the positive samples (Figure 5).

**Figure 5.**
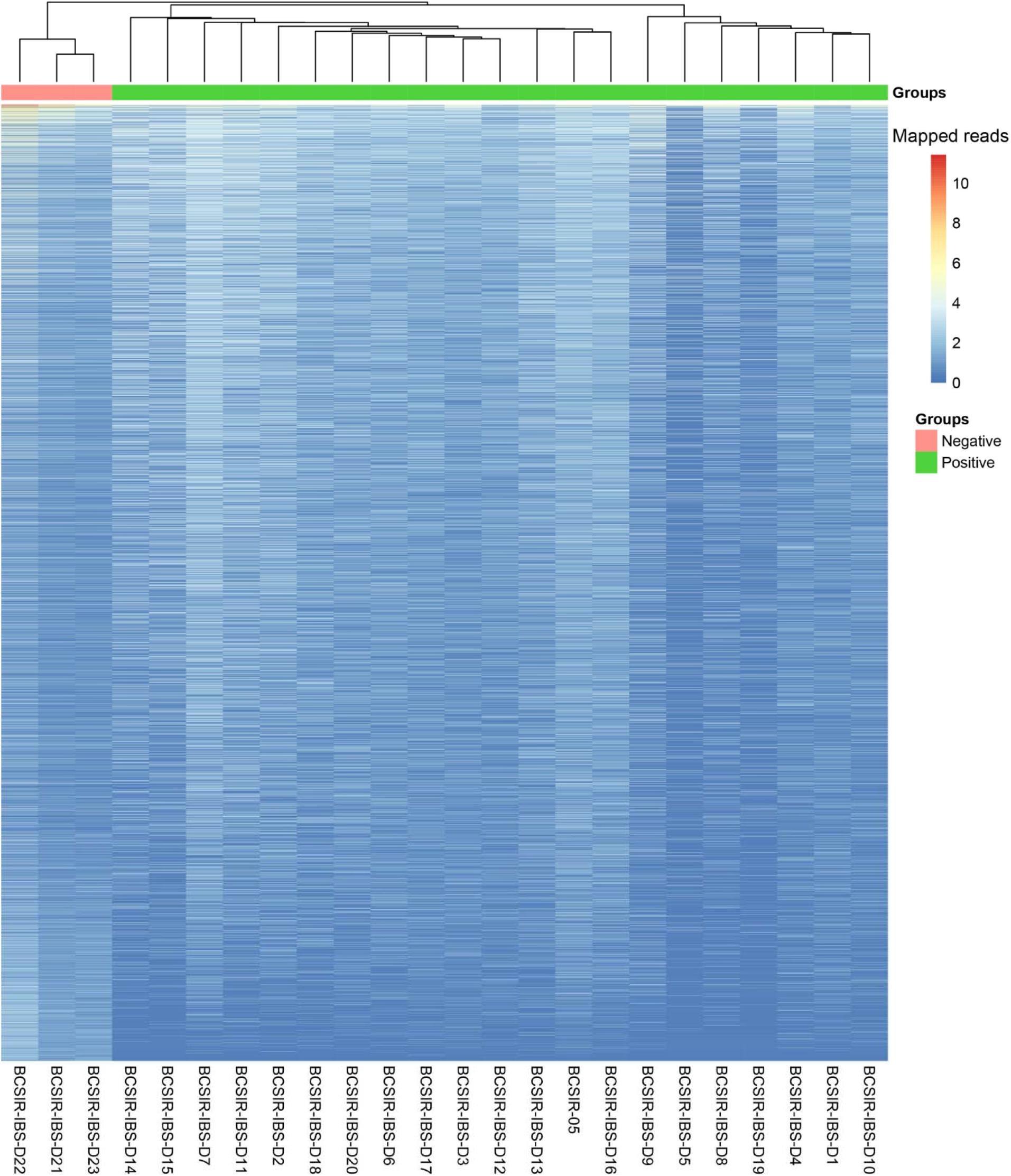
Heatmap of dengue positive and negative cases transcriptome. The genes were analyzed using hierarchical clustering and heatmap, with < 25 genes filtered off every row. In the heatmap, positive and negative groupings create distinct clusters.

Volcano plot represents the DEGs under consideration of default fold change (FC) (log2FC) cut-off is >|2|, and the default p-value cut-off is 0.01 (Figure 6). A total of 11,286 and 1,792 genes exhibited significant fold change (log2FC >|2|) and significant p-value (≤ 0.01), respectively. So the total of (n=1792) genes found as significant DEGs under consideration for both p-values and fold change (FC) (log2FC) (Figure 6, Supplementary data 1). Top three DEGs were identified at positive group over control group: Interleukin 8 (IL-8 (CXCL8) (log2FoldChange = 9.4, p-value = 9.5×10^−08^), X Inactive-Specific Transcript (XIST) log2FoldChange = 7.9, p-value= 1.2×10^−07^), Fucosyltransferase 9 (FUT9) (log2FoldChange 7.9, p-value= 6.9×10-08), these phenomena indicate that the DEGs are associated with dengue (Figure 6).

**Figure 6.**
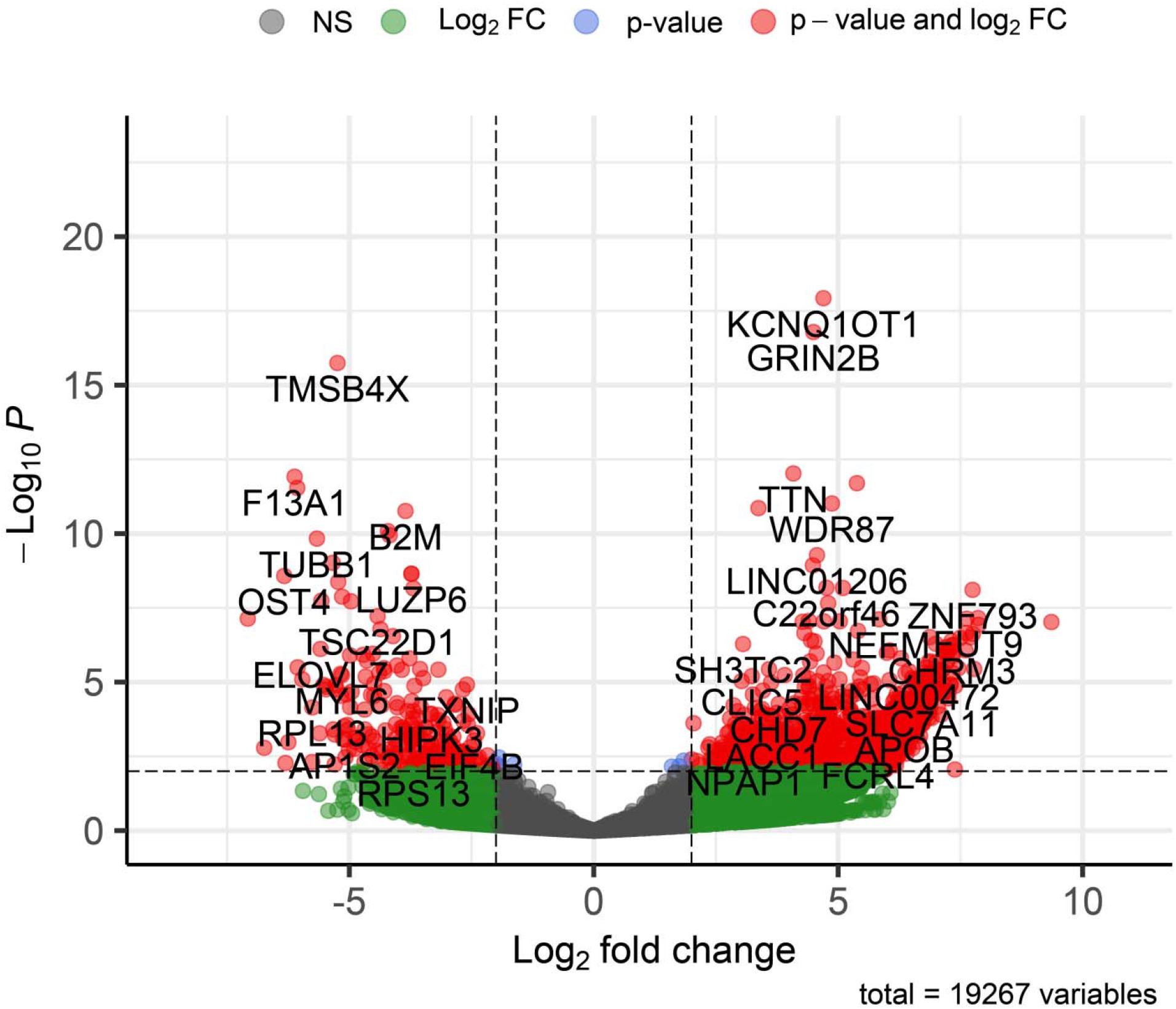
Volcano plots of differentially expressed genes (DEGs) on DEseq2 analysis. The default log_2_FC cut-off is >|2|, and the default p-value cut-off is 0.01. The ash color represents non-significant genes; the green color represents genes that are outside the range of log2FC is −2 to 2 but statistically non-significant (p-value ≥ 0.01); the blue color represents genes that are statistically significant (p-value < 0.01) but outside the range of log2FC is −2 to 2; and the red color represents genes that are statistically and log2FC significant. NS: Non-Significant; FC: Fold Change.

The plot counts function in DESeq2 was used to visualize the top 24 genes with the lowest q-values (Fig. 6, supplementary data 1). The genes include KCNQ1OT1, GRIN2B, TMSB4X, TTN, F13A1, TSIX, PPBP, WDR87, IGFN1, B2M, YWHAZ, MBNL1, TUBB1, LINC01206, NCOA4, LOC440300, LUZP6, MTPN, OST4, RSU1, C22orf46, MEG3, NAP1L1 and ZNF793. Among them, compared to the negative samples, the genes KCNQ1OT1, GRIN2B, TTN, TSIX, WDR87, IGFN1, LINC01206, LOC440300, C22orf46, MEG3, and ZNF793 were upregulated in dengue-positive samples, whereas the others downregulated (Figure 7).

**Figure 7.**
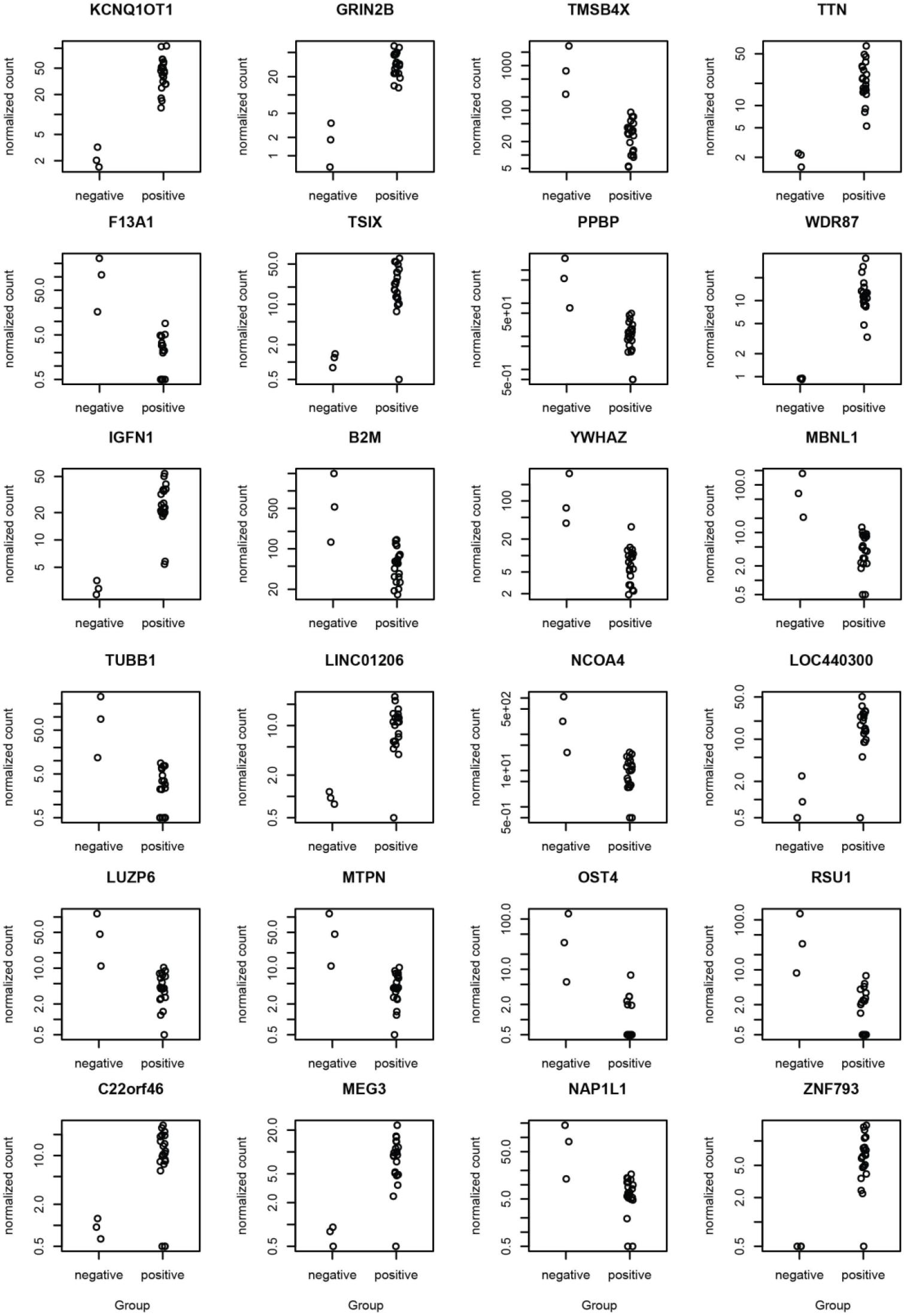
The top 24 DEGs based on p-value adjusted (padj 2.48E-06) were displayed. The Y-axis shows the normalized gene counts, while the X-axis shows the groupings (negative and positive). Details of the analysis can be found in Supplementary Data 1.

### 3.5 Gene Ontology (GO) and pathway analysis of DEGs

To characterize the GO terms, including biological processes, cellular components, molecular functions, and functional pathways of DEGs, we performed over-representation tests in PANTHER version 11.1 (Figure 8 and Supplementary Data 2). We used the GO-Slim PANTHER annotation data set, which represents phylogenetically inferred annotations (Mi et al., 2016).

**Figure 8.**
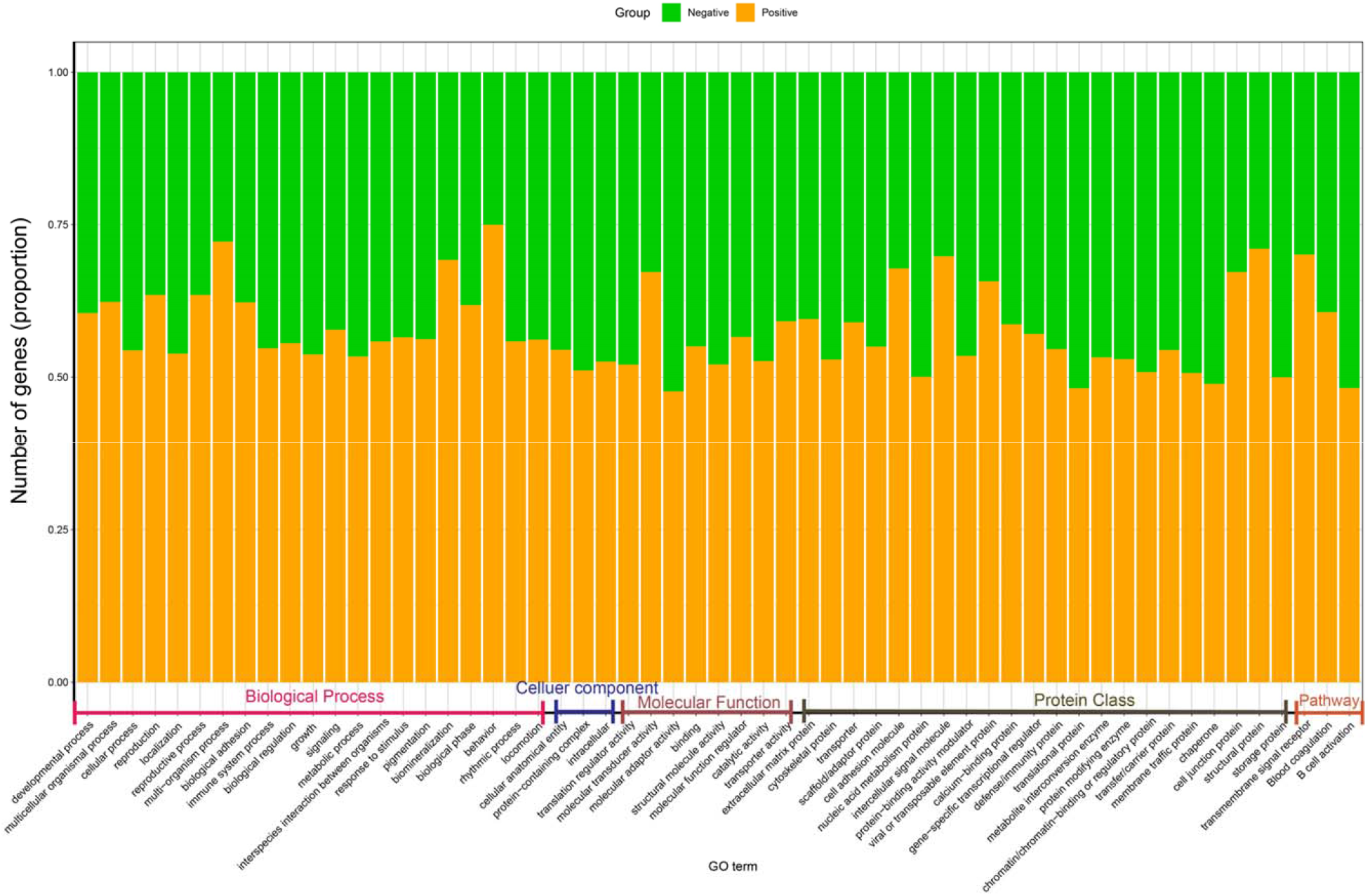
Pathway enrichment and GO phrase analysis for negative and positive groups found in PANTHER Functional classification viewed in gene list analyses. The X-axis of the stack plot shows the GO term, and the Y-axis depicts the proportions of genes for each GO term in each group. The positive group is represented by orange, while the negative group is represented by green. Details of the analysis can be found in Supplementary Data 2.

The current study found overlapping GO terms among comparisons. Interestingly, the olfactory receptor activity (GO:0004984) was identified as the top upregulated (fold change 9.580), and mitochondrial protein complex (GO:0098798) was the top downregulated (fold change 0.640) within the positive group. Within the biological process, the most characterized GO category, compared to the healthy controls, multicellular organismal process (GO:0032501) (fold enrichment 1.32), nervous system process (GO:0050877) (fold enrichment 2.610), system process (GO:0003008) (fold enrichment 2.1), sensory perception of chemical stimulus (GO:0007606) (fold enrichment 4.100) and G protein-coupled receptor signalling pathway (GO:0007186) (fold enrichment 1.610) were significantly upregulated (*p* = < 0.001) in the dengue patients. However, within cellular component, cellular anatomical entity (GO:0110165) (fold change 0.960), protein-containing complex (GO:0032991) (fold change 0.830), intracellular (GO:0005622) (fold change 0.830) significantly decreased (*p* = < 0.001) in dengue patients. Similarly, within category cellular metabolic process (GO:0044237), cellular process (GO:0009987) (fold enrichment 0.950), biological process (GO:0008150) (fold enrichment 0.880), cellular protein metabolic process (GO:0044267) (fold enrichment 0.910) family were significantly downregulated in all the positive samples (Figure 8, supplementary data 2).

Overall, defense/immunity protein class increased in patients (n=166) compared to the healthy controls (n=138), with increased GO of immune system process (GO:0002376) (patients: 329, healthy: 272). Cytokine activity (GO:0005125) significantly (*p* = < 0.001) increased (fold change 2.250) in the patients. However, number of genes decreased related to B cell activation (P00010) (patients: 54, healthy: 58) and T cell activation (P00053) (patients: 72, healthy: 79) (Figure 7, supplementary data 2). Moreover, some GOs are upregulated or downregulated in positive samples such as reproductive process (GO:0022414) (patients/healthy: 120/69), multi-organism process multicellular organismal process (GO:0032501) (1015/613) biological adhesion (GO:0022610) (269/163), blood coagulation (P00011) (37/24), toll receptor signalling pathway (P00054) (53/51), and inflammatory response mediated by chemokine and cytokine signalling pathway (P00031) (196/180) (Supplementary Data 2). Transforming growth factor-beta 1 (TGF-β1) was a highly activated upstream regulator in the positive samples (n=81) compared to the control group (n=60) (Supplementary data 2).

## 4. Discussion

Dengue virus (DENV) is one of the leading threats to public health causing nearly 390 million infections every year worldwide (Tian et al., 2017), with an estimated 2.5 billion people at risk of dengue infection. This mosquito-borne viral disease affects more than 100 endemic countries globally with most disease burden limited to tropical and subtropical regions (Luz et al., 2011). The recent upsurge in the global occurrence of dengue epidemics has prompted research into dengue virus biology and pathology to develop therapeutics for infection prevention and control. Despite numerous efforts to unravel the DENV pathogenesis, the underlying mechanisms of this disease remain elusive and must be investigated further. In this study, we depict the first comprehensive picture of complete genomic characteristics as well as differentially expressed genes in dengue patients in Bangladesh. The expression levels may be implicated to predict the disease progression and, importantly, to design prophylactic therapeutics for the disease.

Dengue infections occur with four distinct serotypes namely DENV1, DENV2, DENV3 and DENV4, posing a major health threat globally. Although mostly the primary infection causes mild to asymptomatic infection with activation of immune responses against DENV serotypes, the severity of the disease is enhanced via heterotypic infection by various serotypes due to antibody-dependent enhancement (ADE) (Roy & Bhattacharjee, 2021). Dengue poses a significant health and economic burden in the WHO South-East Asia Region. Dengue virus has now become hyper endemic in this region for multiple serotypes and genotypes (Tsheten et al., 2021). Co-circulation of all four DENV serotypes is causing the coinfection of these countries with multiple serotypes due to shifts in the dominant serotype (Tsheten et al., 2021). Notably, shift of dominant serotypes from DENV-2 in 2013 to DENV-3 in 2016 occurred in India. Moreover, In 2017, studies conducted in the country revealed that DENV-2 and 3 were dominant in the southern region, while DENV-1 and 2 were prevalent in the northern region (Agarwal et al., 2019; Ganeshkumar et al., 2018; Gupta & Ballani, 2014). Recently, the shifting of predominant serotypes also observed in Thailand (from DENV-2 and DENV-4 to DENV-3), (Hamel et al., 2019) Sri Lanka (from DENV-1 to DENV-2) (Ali et al., 2018), Myanmar (from predominant serotype 1 to co-dominant serotypes 1, 2, and 4) (Oo et al., 2017), Indonesia, Bhutan (from DENV-3 to DENV-1), and in Nepal (Kusmintarsih et al., 2018; Prajapati et al., 2020; Zangmo et al., 2020).

In Bangladesh, DENV-3 of genotype II was predominant till around 2000, which is closely related to the strains from Thailand and Myanmar (M. A. Islam et al., 2006; Podder et al., 2006). From 2013 to 2016, DENV-2 was predominant, with the coexistence of DENV-1. Neither DENV-3 nor DENV-4 were detected during this period (Q. T. Islam, 2019; Muraduzzaman et al., 2018). However, genomic characterization of the outbreak in 2019 having 112000 cases and 129 deaths revealed that DENV type 3 genotype I was predominant, accounting for 93% of tested samples followed by DENV-3 genotype III were detected in two samples from separate districts, and only one DENV-2 cosmopolitan genotype in the Dhaka, capital city of Bangladesh (Titir et al., 2021). Similarly, the present study revealed the predominance of DENV-3 genotype I in all the dengue positive samples (n = 21) from the outbreak of 2021. Interestingly, the viruses closely resemble the strains from China and Thailand. These findings indicate the shift of predominant serotypes from DENV-2 to DENV-3 in the recent outbreaks. The viruses were likely introduced from China or Thailand as previous reports of introduction of the DENV-2 Cosmopolitan genotype from India (Suzuki et al., 2019). These all suggest the continuous cross-border movement of the virus into the South-East Asian region and a requirement for coordinated monitoring and surveillance of DENV (Rico-Hesse et al., 1997; Tsheten et al., 2021). Additionally, these epidemic cycles are potentially responsible for shifting in predominant serotypes and genotypes and have often been associated with higher dengue incidence and severe cases in the territory.

There is a lack of a comprehensive spectrum of gene expression profiles that may contribute to the setting up of dengue severity globally. Several studies have reported gene expression and molecular responses in dengue patients (Devignot et al., 2010; Hoang et al., 2010; Loke et al., 2010; van de Weg et al., 2015). Most of these data produced through the microarray technology lack of a comprehensive spectrum of gene expression patterns due to limited accuracy of expression measurements, especially for transcripts with low abundance (Banerjee et al., 2017). However, recently, the RNA-Seq technology presents a useful tool in disease study with more accuracy at detecting low abundance transcripts and identifying genetic variants (Yu et al., 2019). RNA-Seq also has a wider dynamic range covering the identification of more differentially expressed genes with a higher fold-change in transcript levels of a host (Banerjee et al., 2017). Thus, in the present study, using the RNA-Seq based platform, we provide the first comprehensive overview of the differentially expressed genes in dengue patients compared to healthy controls in Bangladesh.

Our results demonstrated that more genes are expressed in dengue positive samples (n = 17,375) compared to the healthy controls (n = 12,314) with a count of 6,953 and 1,892 unique genes, respectively. These indicate the higher physiological activities stimulated by a viral infection in the patients (Devignot et al., 2010; Hoang et al., 2010). Identification of 2,686 differentially expressed genes (DEGs), with a q-value < 0.05, refers significant alterations in gene expression in dengue patients. Moreover, the heatmap shows distinct clusters of gene expression patterns between patients and healthy individuals (Figure 5). Though the positive samples were subclustered into two groups, no correlation were observed between age, sex and gene expression pattern, thus indicating a similar host response independent of the factors (Hoang et al., 2010). Further, the plot counts function in DESeq2 identified the top 24 genes with the smallest q-values which include KCNQ1OT1, GRIN2B, TMSB4X, TTN, F13A1, TSIX, PPBP, WDR87, IGFN1, B2M, YWHAZ, MBNL1, TUBB1, LINC01206, NCOA4, LOC440300, LUZP6, MTPN, OST4, RSU1, C22orf46, MEG3, NAP1L1 and ZNF793. Among them, compared to the negative samples, the genes KCNQ1OT1, GRIN2B, TTN, TSIX, WDR87, IGFN1, LINC01206, LOC440300, C22orf46, MEG3, and ZNF793 were upregulated in dengue-positive samples, whereas the others downregulated. These genes can be targeted for diagnostic and therapeutic interventions for dengue.

Within the biological process, the most characterized GO category, significant upregulation (*p* = < 0.001) of multicellular organismal process, nervous system process, sensory perception of the chemical stimulus, and G protein-coupled receptor signalling pathway in the dengue patients correlates with the stimulation of various physiological pathways in the patients (Hanley et al., 2021). However, within the cellular component, cellular anatomical entity, protein-containing complex and intracellular significantly decreased (*p* = < 0.001) in dengue patients. Cellular metabolic process, cellular process, biological process and cellular protein metabolic process family were significantly downregulated in all the positive samples, which was contrasting with the previous report (Loke et al., 2010). Importantly, overall, defense/immunity protein class increased in patients compared to the healthy controls, with increased GO of immune system process. Cytokine activity significantly (*p* = < 0.001) increased (fold change 2.250) in the patients. These indicate post-infection immune stimulation in dengue patients, which corroborated with previous findings (Devignot et al., 2010; Loke et al., 2010; van de Weg et al., 2015). Strikingly, number of genes decreased related to B cell activation (patients: 54, healthy: 58) and T cell activation (patients: 72, healthy: 79). To investigate these phenomena, comprehensive studies are warranted aiming at different time points of the infection. Moreover, upregulation of blood coagulation, toll receptor signalling pathway, and inflammation mediated by chemokine and cytokine signalling pathway represent the innate response to the infection (Devignot et al., 2010). Transforming growth factor-beta 1 (TGF-β1) was highly activated upstream regulator in the positive samples could be associated with apoptosis of the infected cells. These differentially expressed genes could be further investigated for target based prophylactic interventions for dengue.

## 5. Conclusion

Dengue remained a major public health threat due to consistent increase in the cases and deaths of infections in Bangladesh. DENV type 3 genotype I is associated with the recent dengue outbreaks in the country. The circulating virus strains closely resemble with the strains from China and Thailand, indicating possible introduction of the DENV-3 genotype I from these countries. Moreover, our results highlight the differentially expressed signature genes associated with dengue, which could be implicated for diagnostic and prophylactic interventions for dengue. Our study has a limitation of small sample size, and this limited our statistical power and our ability to refine validate, and formally compare genetic classifiers. However, we depict the first comprehensive picture of complete genomic characteristics as well as differentially expressed genes in dengue patients in Bangladesh. Further investigation and continuous genomic surveillance are warranted to explore the shift in predominant genotypes and viral pathogenesis.

## Data Availability

Raw sequencing reads generated from this study are deposited at the NCBI sequence read archive (SRA) under accession BioProject PRJNA895688. All the genome accession are available in Table 1.

## Author contributions

MMHS conceptualized the study, collected the samples, processed, run sequencing and finalized the manuscript. MSR analyzed and visualized the data, drafted the manuscript. MRI drafted and critically reviewed the manuscript. AR and MSR analyzed the phylodynamics and phylogenetics of this study. MSI, TAB, SA, GB, IJ, MAH, MMU, MIM, AAS critically reviewed and finalized the manuscript. MSK conceptualized, reviewed and finalized the manuscript.

## Acknowledgement

The authors would like to acknowledge all participants who took part in this study.

## Patient consent

Obtained.

## Ethical approval

The Research Ethics Committee, of Bangladesh Council of Scientific and Industrial Research (BCSIR) approve the research.

---

**Supplementary Figure S1.**
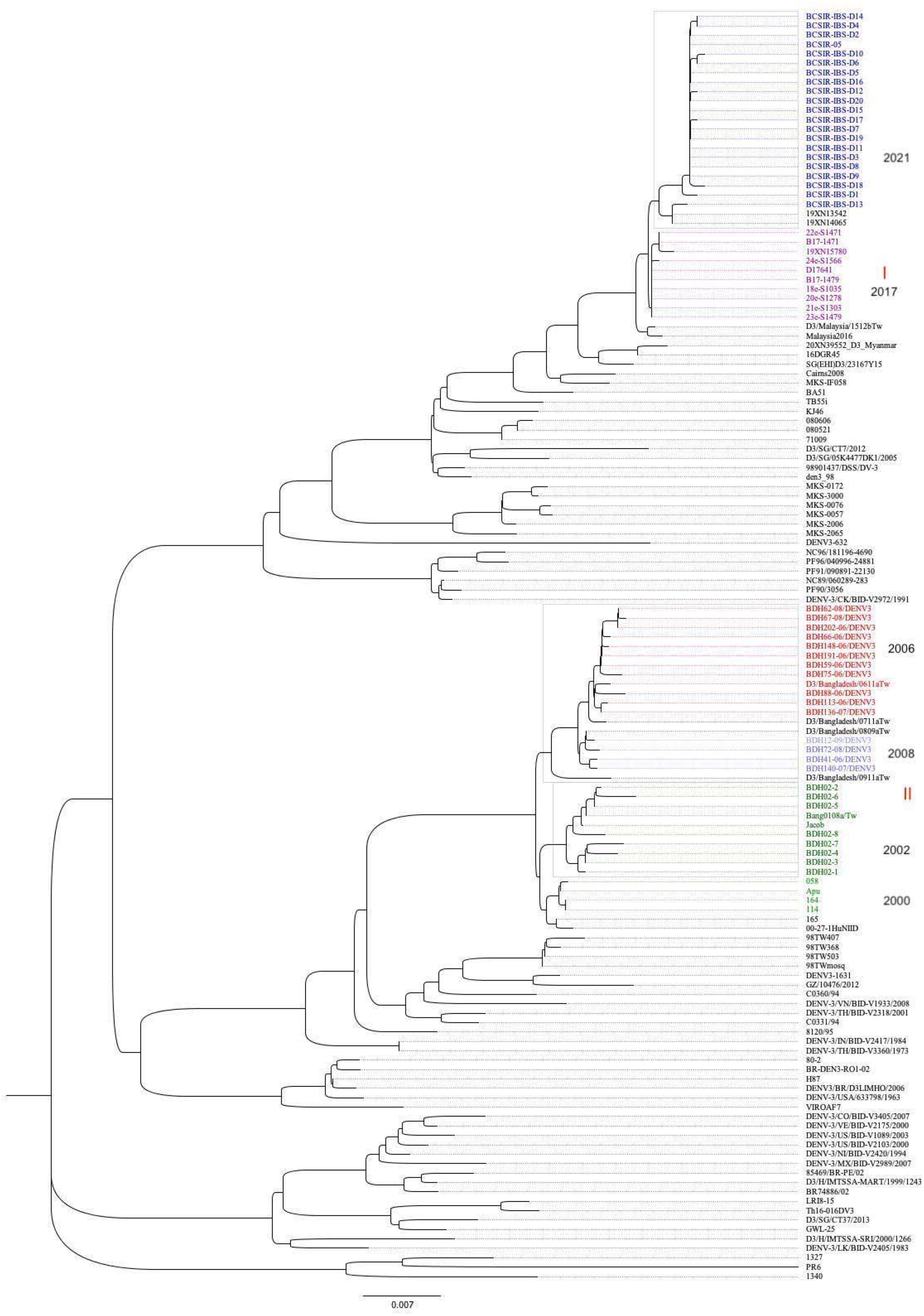
Phylogeny of *env* gene highlighting DENV3 epidemics in Bangladesh. Isolates from all previously reported DENV3 in Bangladesh highlighted in contrasting color along with the year. The 2021 DENV3 epidemic is consistent with previously reported clad switch in 2017 from genotype II to genotype I.

## Reference

Agarwal, A., Gupta, S., Chincholkar, T., Singh, V., Umare, I. K., Ansari, K., Paliya, S., Yadav, A. K., Chowdhary, R., Purwar, S., & Biswas, D. (2019). Co-circulation of dengue virus serotypes in Central India: Evidence of prolonged viremia in DENV-2. Infection, Genetics and Evolution, 70(February), 72–79. https://doi.org/10.1016/j.meegid.2019.02.024

Ali, S., Khan, A. W., Taylor-Robinson, A. W., Adnan, M., Malik, S., & Gul, S. (2018). The unprecedented magnitude of the 2017 dengue outbreak in Sri Lanka provides lessons for future mosquito-borne infection control and prevention. Infection, Disease and Health, 23(2), 114–120. https://doi.org/10.1016/j.idh.2018.02.004

Anders, S., Pyl, P. T., & Huber, W. (2015). HTSeq-A Python framework to work with high-throughput sequencing data. Bioinformatics, 31(2), 166–169. https://doi.org/10.1093/bioinformatics/btu638

Andrews, S., & others. (2010). FastQC: a quality control tool for high throughput sequence data. 2010. In https://www.bioinformatics.babraham.ac.uk/projects/fastqc/ (p. http://www.bioinformatics.babraham.ac.uk/projects/).

Aziz, M. M., Hasan, K. N., Hasanat, M. A., Siddiqui, M. A., Salimullah, M., Chowdhury, A. K., Ahmed, M., Alam, M. N., & Hassan, M. S. (2002). Predominance of the DEN-3 genotype during the recent dengue outbreak in Bangladesh. The Southeast Asian Journal of Tropical Medicine and Public Health, 33(1), 42–48.

Banerjee, A., Shukla, S., Pandey, A. D., Goswami, S., Bandyopadhyay, B., Ramachandran, V., Das, S., Malhotra, A., Agarwal, A., Adhikari, S., Rahman, M., Chatterjee, S., Bhattacharya, N., Basu, N., Pandey, P., Sood, V., & Vrati, S. (2017). RNA-Seq analysis of peripheral blood mononuclear cells reveals unique transcriptional signatures associated with disease progression in dengue patients. Translational Research, 186, 62–78.e9. https://doi.org/10.1016/j.trsl.2017.06.007

Benjamini, Y., & Hochberg, Y. (1995). Controlling the False Discovery Rate: A Practical and Powerful Approach to Multiple Testing. Journal of the Royal Statistical Society: Series B (Methodological), 57(1), 289–300. https://doi.org/10.1111/j.2517-6161.1995.tb02031.x

Bhatt, S., Gething, P. W., Brady, O. J., Messina, J. P., Farlow, A. W., Moyes, C. L., Drake, J. M., Brownstein, J. S., Hoen, A. G., Sankoh, O., Myers, M. F., George, D. B., Jaenisch, T., William Wint, G. R., Simmons, C. P., Scott, T. W., Farrar, J. J., & Hay, S. I. (2013). The global distribution and burden of dengue. Nature, 496(7446), 504–507. https://doi.org/10.1038/nature12060

Bolger, A. M., Lohse, M., & Usadel, B. (2014). Trimmomatic: A flexible trimmer for Illumina sequence data. Bioinformatics, 30(15), 2114–2120. https://doi.org/10.1093/bioinformatics/btu170

Boni, M. F., Posada, D., & Feldman, M. W. (2007). An exact nonparametric method for inferring mosaic structure in sequence triplets. Genetics, 176(2), 1035–1047. https://doi.org/10.1534/genetics.106.068874

Bournazos, S., Gupta, A., & Ravetch, J. V. (2020). The role of IgG Fc receptors in antibody-dependent enhancement. Nature Reviews Immunology, 20(10), 633–643. https://doi.org/10.1038/s41577-020-00410-0

Buchfink, B., Xie, C., & Huson, D. H. (2014). Fast and sensitive protein alignment using DIAMOND. Nature Methods, 12(1), 59–60. https://doi.org/10.1038/nmeth.3176

Carpenter, S., Aiello, D., Atianand, M. K., Ricci, E. P., Gandhi, P., Hall, L. L., Byron, M., Monks, B., Henry-Bezy, M., Lawrence, J. B., O’Neill, L. A. J., Moore, M. J., Caffrey, D. R., & Fitzgerald, K. A. (2013). A long noncoding RNA mediates both activation and repression of immune response genes. Science, 341(6147), 789–792. https://doi.org/10.1126/science.1240925

Chareonsirisuthigul, T., Kalayanarooj, S., & Ubol, S. (2007). Dengue virus (DENV) antibody-dependent enhancement of infection upregulates the production of anti-inflammatory cytokines, but suppresses anti-DENV free radical and pro-inflammatory cytokine production, in THP-1 cells. Journal of General Virology, 88(2), 365–375. https://doi.org/10.1099/vir.0.82537-0

Chu, C. P., Hokamp, J. A., Cianciolo, R. E., Dabney, A. R., Brinkmeyer-Langford, C., Lees, G. E., & Nabity, M. B. (2017). RNA-seq of serial kidney biopsies obtained during progression of chronic kidney disease from dogs with X-linked hereditary nephropathy. Scientific Reports, 7(1), 1–14. https://doi.org/10.1038/s41598-017-16603-y

Deforche, K. (2017). An alignment method for nucleic acid sequences against annotated genomes. BioRxiv, 1–15.

Deng, S. Q., Yang, X., Wei, Y., Chen, J. T., Wang, X. J., & Peng, H. J. (2020). A review on dengue vaccine development. Vaccines, 8(1), 1–13. https://doi.org/10.3390/vaccines8010063

Devignot, S., Sapet, C., Duong, V., Bergon, A., Rihet, P., Ong, S., Lorn, P. T., Chroeung, N., Ngeav, S., Tolou, H. J., Buchy, P., & Couissinier-Paris, P. (2010). Genome-wide expression profiling deciphers host responses altered during dengue shock syndrome and reveals the role of innate immunity in severe dengue. PLoS ONE, 5(7). https://doi.org/10.1371/journal.pone.0011671

Dobin, A., Davis, C. A., Schlesinger, F., Drenkow, J., Zaleski, C., Jha, S., Batut, P., Chaisson, M., & Gingeras, T. R. (2013). STAR: Ultrafast universal RNA-seq aligner. Bioinformatics, 29(1), 15–21. https://doi.org/10.1093/bioinformatics/bts635

Drummond, A. J., & Rambaut, A. (2007). BEAST: Bayesian evolutionary analysis by sampling trees. BMC Evolutionary Biology, 7(1), 1–8. https://doi.org/10.1186/1471-2148-7-214

Erickson, K. (2010). The jukes-cantor model of molecular evolution. Primus, 20(5), 438–445. https://doi.org/10.1080/10511970903487705

Ganeshkumar, P., Murhekar, M. V., Poornima, V., Saravanakumar, V., Sukumaran, K., Anandaselvasankar, A., John, D., & Mehendale, S. M. (2018). Dengue infection in India: A systematic review and meta-analysis. PLoS Neglected Tropical Diseases, 12(7), 2–3. https://doi.org/10.1371/journal.pntd.0006618

Gibbs, M. J., Armstrong, J. S., & Gibbs, A. J. (2000). Sister-scanning: A Monte Carlo procedure for assessing signals in rebombinant sequences. Bioinformatics, 16(7), 573–582. https://doi.org/10.1093/bioinformatics/16.7.573

Gupta, E., & Ballani, N. (2014). Current perspectives on the spread of dengue in India. Infection and Drug Resistance, 7, 337–342. https://doi.org/10.2147/IDR.S55376

Hamel, R., Surasombatpattana, P., Wichit, S., Dauvé, A., Donato, C., Pompon, J., Vijaykrishna, D., Liegeois, F., Vargas, R. M., Luplertlop, N., & Missé, D. (2019). Phylogenetic analysis revealed the co-circulation of four dengue virus serotypes in Southern Thailand. PLoS ONE, 14(8), 1–16. https://doi.org/10.1371/journal.pone.0221179

Hanley, J. P., Tu, H. A., Dragon, J. A., Dickson, D. M., Rio-Guerra, R. del, Tighe, S. W., Eckstrom, K. M., Selig, N., Scarpino, S. V., Whitehead, S. S., Durbin, A. P., Pierce, K. K., Kirkpatrick, B. D., Rizzo, D. M., Frietze, S., & Diehl, S. A. (2021). Immunotranscriptomic profiling the acute and clearance phases of a human challenge dengue virus serotype 2 infection model. Nature Communications, 12(1), 1–14. https://doi.org/10.1038/s41467-021-22930-6

Ho, S. Y. W., & Shapiro, B. (2011). Skyline-plot methods for estimating demographic history from nucleotide sequences. Molecular Ecology Resources, 11(3), 423–434. https://doi.org/10.1111/j.1755-0998.2011.02988.x

Hoang, L. T., Lynn, D. J., Henn, M., Birren, B. W., Lennon, N. J., Le, P. T., Duong, K. T. H., Nguyen, T. T. H., Mai, L. N., Farrar, J. J., Hibberd, M. L., & Simmons, C. P. (2010). The Early Whole-Blood Transcriptional Signature of Dengue Virus and Features Associated with Progression to Dengue Shock Syndrome in Vietnamese Children and Young Adults. Journal of Virology, 84(24), 12982–12994. https://doi.org/10.1128/jvi.01224-10

Holmes, E. C., & Twiddy, S. S. (2003). The origin, emergence and evolutionary genetics of dengue virus. Infection, Genetics and Evolution, 3(1), 19–28. https://doi.org/10.1016/S1567-1348(03)00004-2

Hur, S. (2019). Double-Stranded RNA Sensors and Modulators in Innate Immunity. Annual Review of Immunology, 37, 349–375. https://doi.org/10.1146/annurev-immunol-042718-041356

Islam, M. A., Ahmed, M. U., Begum, N., Chowdhury, N. A., Khan, A. H., Parquet, M. del C., Bipolo, S., Inoue, S., Hasebe, F., Suzuki, Y., & Morita, K. (2006). Molecular characterization and clinical evaluation of dengue outbreak in 2002 in Bangladesh. Japanese Journal of Infectious Diseases, 59(2), 85–91.

Islam, Q. T. (2019). Changing epidemiological and clinical pattern of dengue in bangladesh 2018. Journal of Medicine (Bangladesh), 20(1), 1–3. https://doi.org/10.3329/jom.v20i1.38812

Iwasaki, A., & Medzhitov, R. (2015). Control of adaptive immunity by the innate immune system. Nature Immunology, 16(4), 343–353. https://doi.org/10.1038/ni.3123

Katoh, K., & Standley, D. M. (2013). MAFFT multiple sequence alignment software version 7: Improvements in performance and usability. Molecular Biology and Evolution, 30(4), 772–780. https://doi.org/10.1093/molbev/mst010

Kobasa, D., Jones, S. M., Shinya, K., Kash, J. C., Copps, J., Ebihara, H., Hatta, Y., Kim, J. H., Halfmann, P., Hatta, M., Feldmann, F., Alimonti, J. B., Fernando, L., Li, Y., Katze, M. G., Feldmann, H., & Kawaoka, Y. (2007). Aberrant innate immune response in lethal infection of macaques with the 1918 influenza virus. Nature, 445(7125), 319–323. https://doi.org/10.1038/nature05495

Kolde, R. (2015). pheatmap◻: Pretty Heatmaps. R Package Version 1.0.8, 1–7. https://cran.r-project.org/web/packages/pheatmap/pheatmap.pdf

Kumar, S., Stecher, G., Peterson, D., & Tamura, K. (2012). MEGA-CC: Computing core of molecular evolutionary genetics analysis program for automated and iterative data analysis. Bioinformatics, 28(20), 2685–2686. https://doi.org/10.1093/bioinformatics/bts507

Kusmintarsih, E. S., Hayati, R. F., Turnip, O. N., Yohan, B., Suryaningsih, S., Pratiknyo, H., Denis, D., & Sasmono, R. T. (2018). Molecular characterization of dengue viruses isolated from patients in Central Java, Indonesia. Journal of Infection and Public Health, 11(5), 617–625. https://doi.org/10.1016/j.jiph.2017.09.019

Lazear, H. M., Schoggins, J. W., & Diamond, M. S. (2019). Shared and Distinct Functions of Type I and Type III Interferons. Immunity, 50(4), 907–923. https://doi.org/10.1016/j.immuni.2019.03.025

Li, H., & Durbin, R. (2009). Fast and accurate short read alignment with Burrows-Wheeler transform. Bioinformatics, 25(14), 1754–1760. https://doi.org/10.1093/bioinformatics/btp324

Li, H., Handsaker, B., Wysoker, A., Fennell, T., Ruan, J., Homer, N., Marth, G., Abecasis, G., & Durbin, R. (2009). The Sequence Alignment/Map format and SAMtools. Bioinformatics, 25(16), 2078–2079. https://doi.org/10.1093/bioinformatics/btp352

Li, M. J., Lan, C. J., Gao, H. T., Xing, D., Gu, Z. Y., Su, D., Zhao, T. Y., Yang, H. Y., & Li, C. X. (2020). Transcriptome analysis of Aedes aegypti Aag2 cells in response to dengue virus-2 infection. Parasites and Vectors, 13(1), 1–14. https://doi.org/10.1186/s13071-020-04294-w

Loke, P., Hammond, S. N., Leung, J. M., Kim, C. C., Batra, S., Rocha, C., Balmaseda, A., & Harris, E. (2010). Gene expression patterns of dengue virus-infected children from nicaragua reveal a distinct signature of increased metabolism. PLoS Neglected Tropical Diseases, 4(6). https://doi.org/10.1371/journal.pntd.0000710

Love, M. I., Huber, W., & Anders, S. (2014). Moderated estimation of fold change and dispersion for RNA-seq data with DESeq2. Genome Biology, 15(12), 1–21. https://doi.org/10.1186/s13059-014-0550-8

Luz, P. M., Vanni, T., Medlock, J., Paltiel, A. D., & Galvani, A. P. (2011). Dengue vector control strategies in an urban setting: An economic modelling assessment. The Lancet, 377(9778), 1673–1680. https://doi.org/10.1016/S0140-6736(11)60246-8

Mahbubur Rahman, K. R., A. K. Siddque, Shereen Shoma, A. H. M. K., & K. S. Ali, Ananda Nisaluk, and R. F. B. (2002). First Outbreak of. Emerging Infectious Diseases, 8(7), 2000–2002.

Martin, D. P., Posada, D., Crandall, K. A., & Williamson, C. (2005). A modified bootscan algorithm for automated identification of recombinant sequences and recombination breakpoints. AIDS Research and Human Retroviruses, 21(1), 98–102. https://doi.org/10.1089/aid.2005.21.98

Martin, Darren P., Murrell, B., Golden, M., Khoosal, A., & Muhire, B. (2015). RDP4: Detection and analysis of recombination patterns in virus genomes. Virus Evolution, 1(1), 1–5. https://doi.org/10.1093/ve/vev003

Martin, Darren P., Varsani, A., Roumagnac, P., Botha, G., Maslamoney, S., Schwab, T., Kelz, Z., Kumar, V., & Murrell, B. (2021). RDP5: A computer program for analyzing recombination in, and removing signals of recombination from, nucleotide sequence datasets. Virus Evolution, 7(1), 5–7. https://doi.org/10.1093/ve/veaa087

Medzhitov, R. (2007). Recognition of microorganisms and activation of the immune response. Nature, 449(7164), 819–826. https://doi.org/10.1038/nature06246

Mi, H., Ebert, D., Muruganujan, A., Mills, C., Albou, L. P., Mushayamaha, T., & Thomas, P. D. (2021). PANTHER version 16: A revised family classification, tree-based classification tool, enhancer regions and extensive API. Nucleic Acids Research, 49(D1), D394–D403. https://doi.org/10.1093/nar/gkaa1106

Mi, H., Poudel, S., Muruganujan, A., Casagrande, J. T., & Thomas, P. D. (2016). PANTHER version 10: Expanded protein families and functions, and analysis tools. Nucleic Acids Research, 44(D1), D336–D342. https://doi.org/10.1093/nar/gkv1194

Mukhopadhyay, S., Kuhn, R. J., & Rossmann, M. G. (2005). A structural perspective of the Flavivirus life cycle. Nature Reviews Microbiology, 3(1), 13–22. https://doi.org/10.1038/nrmicro1067

Muñoz-Jordán, J. L., Sánchez-Burgos, G. G., Laurent-Rolle, M., & García-Sastre, A. (2003). Inhibition of interferon signaling by dengue virus. Proceedings of the National Academy of Sciences of the United States of America, 100(SUPPL. 2), 14333–14338. https://doi.org/10.1073/pnas.2335168100

Muraduzzaman, A. K. M., Alam, A. N., Sultana, S., Siddiqua, M., Khan, M. H., Akram, A., Haque, F., Flora, M. S., & Shirin, T. (2018). Circulating dengue virus serotypes in Bangladesh from 2013 to 2016. VirusDisease, 29(3), 303–307. https://doi.org/10.1007/s13337-018-0469-x

Nikolayeva, I., Bost, P., Casademont, I., Duong, V., Koeth, F., Prot, M., Czerwinska, U., Ly, S., Bleakley, K., Cantaert, T., Dussart, P., Buchy, P., Simon-Lorière, E., Sakuntabhai, A., & Schwikowski, B. (2018). A blood RNA signature detecting severe disease in young dengue patients at hospital arrival. Journal of Infectious Diseases, 217(11), 1690–1698. https://doi.org/10.1093/infdis/jiy086

Nunes, P. C. G., Sampaio, S. A. F., Rodrigues da Costa, N., de Mendonça, M. C. L., Lima, M. da R. Q., Araujo, S. E. M., dos Santos, F. B., Simões, J. B. S., de Santis Gonçalves, B., Nogueira, R. M. R., & de Filippis, A. M. B. (2016). Dengue severity associated with age and a new lineage of dengue virus-type 2 during an outbreak in Rio De Janeiro, Brazil. Journal of Medical Virology, 88(7), 1130–1136. https://doi.org/10.1002/jmv.24464

Nurk, S., Meleshko, D., Korobeynikov, A., & Pevzner, P. A. (2017). MetaSPAdes: A new versatile metagenomic assembler. Genome Research, 27(5), 824–834. https://doi.org/10.1101/gr.213959.116

Oo, P. M., Wai, K. T., Harries, A. D., Shewade, H. D., Oo, T., Thi, A., & Lin, Z. (2017). The burden of dengue, source reduction measures, and serotype patterns in Myanmar, 2011 to 2015-R2. Tropical Medicine and Health, 45(1), 1–11. https://doi.org/10.1186/s41182-017-0074-5

Padidam, M., Sawyer, S., & Fauquet, C. M. (1999). Possible emergence of new geminiviruses by frequent recombination. Virology, 265(2), 218–225. https://doi.org/10.1006/viro.1999.0056

Patro, R., Duggal, G., Love, M. I., Irizarry, R. A., & Kingsford, C. (2017). Salmon provides fast and bias-aware quantification of transcript expression. Nature Methods, 14(4), 417–419. https://doi.org/10.1038/nmeth.4197

Podder, G., Breiman, R. F., Azim, T., Thu, H. M., Velathanthiri, N., Mai, L. Q., Lowry, K., & Aaskov, J. G. (2006). Origin of dengue type 3 viruses associated with the dengue outbreak in Dhaka, Bangladesh, in 2000 and 2001. American Journal of Tropical Medicine and Hygiene, 74(2), 263–265. https://doi.org/10.4269/ajtmh.2006.74.263

Posada, D., & Crandall, K. A. (2001). Evaluation of methods for detecting recombination from DNA sequences: Computer simulations. Proceedings of the National Academy of Sciences of the United States of America, 98(24), 13757–13762. https://doi.org/10.1073/pnas.241370698

Prajapati, S., Napit, R., Bastola, A., Rauniyar, R., Shrestha, S., Lamsal, M., Adhikari, A., Bhandari, P., Yadav, S. R., & Manandhar, K. Das. (2020). Molecular phylogeny and distribution of dengue virus serotypes circulating in Nepal in 2017. PLoS ONE, 15(7 July), 1–17. https://doi.org/10.1371/journal.pone.0234929

Quinlan, A. R., & Hall, I. M. (2010). BEDTools: A flexible suite of utilities for comparing genomic features. Bioinformatics, 26(6), 841–842. https://doi.org/10.1093/bioinformatics/btq033

Rambaut, A., Drummond, A. J., Xie, D., Baele, G., & Suchard, M. A. (2018). Posterior summarization in Bayesian phylogenetics using Tracer 1.7. Systematic Biology, 67(5), 901–904. https://doi.org/10.1093/sysbio/syy032

Rico-Hesse, R., Harrison, L. M., Salas, R. A., Tovar, D., Nisalak, A., Ramos, C., Boshell, J., De Mesa, M. T. R., Nogueira, R. M. R., & Rosa, A. T. Da. (1997). Origins of dengue type 2 viruses associated with increased pathogenicity in the Americas. Virology, 230(2), 244–251. https://doi.org/10.1006/viro.1997.8504

Roy, S. K., & Bhattacharjee, S. (2021). Dengue virus: Epidemiology, biology, and disease aetiology. Canadian Journal of Microbiology, 67(10), 687–702. https://doi.org/10.1139/cjm-2020-0572

Saini, J., Bandyopadhyay, B., Pandey, A. D., Ramachandran, V. G., Das, S., Sood, V., Banerjee, A., & Vrati, S. (2020). High-Throughput RNA Sequencing Analysis of Plasma Samples Reveals Circulating microRNA Signatures with Biomarker Potential in Dengue Disease Progression. MSystems, 5(5). https://doi.org/10.1128/msystems.00724-20

Saitou, N., & Nei, M. (1987). The neighbor-joining method: a new method for reconstructing phylogenetic trees. Molecular Biology and Evolution, 4(4), 406–425. https://doi.org/10.1093/oxfordjournals.molbev.a040454

Seemann, T. (2015). Snippy-Rapid haploid variant calling and core SNP phylogeny. In GitHub.

Smith, J. M. (1992). Analyzing the mosaic structure of genes. Journal of Molecular Evolution, 34(2), 126–129. https://doi.org/10.1007/BF00182389

Souza-Neto, J. A., Sim, S., & Dimopoulos, G. (2009). An evolutionary conserved function of the JAK-STAT pathway in anti-dengue defense. Proceedings of the National Academy of Sciences of the United States of America, 106(42), 17841–17846. https://doi.org/10.1073/pnas.0905006106

Suzuki, K., Phadungsombat, J., Nakayama, E. E., Saito, A., Egawa, A., Sato, T., Rahim, R., Hasan, A., Lin, M. Y. C., Takasaki, T., Rahman, M., & Shioda, T. (2019). Genotype replacement of dengue virus type 3 and clade replacement of dengue virus type 2 genotype Cosmopolitan in Dhaka, Bangladesh in 2017. Infection, Genetics and Evolution, 75(July). https://doi.org/10.1016/j.meegid.2019.103977

Tian, H., Sun, Z., Faria, N. R., Yang, J., Cazelles, B., Huang, S., Xu, B., Yang, Q., Pybus, O. G., & Xu, B. (2017). Increasing airline travel may facilitate co-circulation of multiple dengue virus serotypes in Asia. PLoS Neglected Tropical Diseases, 11(8), 1–15. https://doi.org/10.1371/journal.pntd.0005694

Titir, S. R., Paul, S. K., Ahmed, S., Haque, N., Nasreen, S. A., Hossain, K. S., Ahmad, F. U., Nila, S. S., Khanam, J., Nowsher, N., Amin, A. M. M. Al, Khan, A. U., Aung, M. S., & Kobayashi, N. (2021). Nationwide distribution of dengue virus type 3 (Denv-3) genotype i and emergence of denv-3 genotype iii during the 2019 outbreak in bangladesh. Tropical Medicine and Infectious Disease, 6(2). https://doi.org/10.3390/tropicalmed6020058

Tsheten, T., Gray, D. J., Clements, A. C. A., & Wangdi, K. (2021). Epidemiology and challenges of dengue surveillance in the WHO South-East Asia Region. Transactions of the Royal Society of Tropical Medicine and Hygiene, 115(6), 583–599. https://doi.org/10.1093/trstmh/traa158

van de Weg, C. A. M., van den Ham, H. J., Bijl, M. A., Anfasa, F., Zaaraoui-Boutahar, F., Dewi, B. E., Nainggolan, L., van IJcken, W. F. J., Osterhaus, A. D. M. E., Martina, B. E. E., van Gorp, E. C. M., & Andeweg, A. C. (2015). Time since Onset of Disease and Individual Clinical Markers Associate with Transcriptional Changes in Uncomplicated Dengue. PLoS Neglected Tropical Diseases, 9(3), 1–20. https://doi.org/10.1371/journal.pntd.0003522

Vilsker, M., Moosa, Y., Nooij, S., Fonseca, V., Ghysens, Y., Dumon, K., Pauwels, R., Alcantara, L. C., Vanden Eynden, E., Vandamme, A. M., Deforche, K., & De Oliveira, T. (2019). Genome Detective: An automated system for virus identification from high-throughput sequencing data. Bioinformatics, 35(5), 871–873. https://doi.org/10.1093/bioinformatics/bty695

Wickham, H. (2011). Ggplot2. Wiley Interdisciplinary Reviews: Computational Statistics, 3(2), 180–185. https://doi.org/10.1002/wics.147

Yu, J., Peterson, D. R., Baran, A. M., Bhattacharya, S., Wylie, T. N., Falsey, A. R., Mariani, T. J., & Storch, G. A. (2019). Host gene expression in nose and blood for the diagnosis of viral respiratory infection. Journal of Infectious Diseases, 219(7), 1151–1161. https://doi.org/10.1093/infdis/jiy608

Zangmo, S., Darnal, J. B., Tsheten, Gyeltshen, S., Thapa, B. T., Rodpradit, P., Chinnawirotpisan, P., Manasatienkij, W., Macareo, L. R., Fernandez, S., Wangchuk, S., & Klungthong, C. (2020). Molecular epidemiology of dengue fever outbreaks in Bhutan, 2016-2017. PLoS Neglected Tropical Diseases, 14(4), 1–12. https://doi.org/10.1371/journal.pntd.0008165

Zeng, Z., Shi, J., Guo, X., Mo, L., Hu, N., Sun, J., Wu, M., Zhou, H., & Hu, Y. (2018). Full-length genome and molecular characterization of dengue virus serotype 2 isolated from an imported patient from Myanmar. Virology Journal, 15(1), 1–12. https://doi.org/10.1186/s12985-018-1043-2

